## Supplementary material for "ArtSymbioCyc, a metabolic network database collection dedicated to arthropod symbioses: a case study, the tripartite cooperation in *Sipha maydis*": Sup FIG S1-S2, Sup Tables S1-S3

**Table S1.** Databases currently contained in ArtSymbioCyc and specific features of the corresponding metabolic networks.

| Holobionts | Accession Numbers (genome reference) | Pathway numbers | Enzyme numbers | Reaction numbers | Compound numbers |
|---|---|---|---|---|---|
| Api-*Acyrthosiphon pisum* holobiont | - | 689 | 4,747 | 4,246 | 2,577 |
| Api-*Acyrthosiphon pisum* holobiont (*Buchnera* only) | - | 619 | 4,243 | 3,937 | 2,352 |
| Api-*Acyrthosiphon pisum* AL4f | GCF_005508785.2 (1) | 289 | 3,897 | 2,536 | 1,447 |
| Api-*Buchnera aphidicola* APS | GCF_000009605.1 (2) | 98 | 329 | 693 | 500 |
| Api-*Candidatus* Hamiltonella defensa T5A | GCF_000021705.1 (3) | 161 | 523 | 1,037 | 749 |
| Sma-*Sipha maydis* holobiont | - | 616 | 3,175 | 3,910 | 2,270 |
| Sma-*Sipha maydis* Midelt | GCA_034509805.1 (4) | 273 | 2,523 | 2,409 | 1,344 |
| Sma-*Buchnera aphidicola* Sm_Midelt | GCF_024029855.1 (5) | 58 | 221 | 473 | 321 |
| Sma-*Serratia symbiotica* Sm_Midelt | GCA_024160085.1 (5) | 135 | 410 | 899 | 617 |
| Cce-*Cinara cedri* holobiont | - | 619 | 3,903 | 3,994 | 2,387 |
| Cce-*Cinara cedri* | GCA_902439185.1 (6) | 274 | 3,426 | 2,458 | 1,488 |
| Cce-*Buchnera* aphidicola BCc | GCF_000090965.1(7, 8) | 41 | 189 | 402 | 320 |
| Cce-Serratia symbiotica Cc | GCA_000238975.1 (9) | 116 | 359 | 754 | 599 |
| Bta_MEAM1-*Bemisia tabaci* MEAM1 holobiont | - | 702 | 3,477 | 4,392 | 2,613 |
| Bta_MEAM1-*Bemisia tabaci* MEAM1 | MEAM1 V1.2 (10) | 295 | 2,442 | 2,584 | 1,491 |
| Bta_MEAM1-*Candidatus* Portiera aleyrodidarum | *Candidatus* Portiera[a] (10) | 33 | 131 | 286 | 213 |
| Bta_MEAM1-*Hamiltonella defensa* | *H. defensa*[a] (11) | 179 | 537 | 1060 | 712 |
| Bta_MEAM1-*Rickettsia sp.* | *Rickettsia sp.*[a] (12) | 117 | 363 | 784 | 534 |

| | | | | | |
|---|---|---|---|---|---|
| Bta_MED-*Bemisia tabaci* MED holobiont | - | 766 | 4,639 | 4,696 | 2,757 |
| Bta_MED-*Bemisia tabaci* MED | MED v1.0[b] (13) | 319 | 3,295 | 2949 | 1762 |
| Bta_MED-*Candidatus* Portiera aleyrodidarum BT-QVLC | GCA_000298385.1 (14) | 36 | 142 | 313 | 232 |
| Bta_MED-*Hamiltonella defensa* | GCF_000258345.2 (15) | 182 | 550 | 1206 | 768 |
| Bta_MED-*Candidatus* Cardinium hertigii | GCF_000689375.1 (16) | 74 | 241 | 669 | 386 |
| Bta_MED-*Wolbachia sp.* | GCF_900097055.1 (17) | 114 | 365 | 861 | 540 |
| Cle-*Cimex lectularius* holobiont | - | 564 | 4,322 | 3,863 | 2,241 |
| Cle-*Cimex lectularius* Harlan | GCF_000648675.2 (18) | 287 | 3,922 | 2,464 | 1,418 |
| Cle-*Wolbachia sp.* | GCF_000829315.1 (19) | 112 | 391 | 880 | 557 |
| Dme-*Drosophila melanogaster* holobiont | - | 783 | 6,402 | 4,466 | 2,715 |
| Dme-*Drosophila melanogaster* holobiont (gut microbiota) | - | 779 | 6,078 | 4,448 | 2,673 |
| Dme-*Drosophila melanogaster* | GCF_000001215.4 (20) | 296 | 3,857 | 2,333 | 1,307 |
| Dme-*Lactiplantibacillus plantarum plantarum NC8* | AGRI01000001.1 (21) | 232 | 846 | 1,379 | 1,001 |
| Dme-*Acetobacter pomorum* WJL DM001 | PRJNA60787 (22) | 286 | 1,241 | 1,612 | 1,179 |
| Dme-*Wolbachia sp.* | GCF_016584425.1 (23) | 108 | 340 | 735 | 549 |
| Gmo-*Glossina morsitans* holobiont | - | 676 | 3,429 | 3,943 | 2,360 |
| - Gmo-*Glossina morsitans Yale* | GmorY1[c] (24) | 284 | 2,258 | 2,273 | 1,370 |
| - Gmo-*Sodalis glossinidius morsitans* | GCF_000010085.1 (25) | 274 | 826 | 1,609 | 1,191 |
| - Gmo-*Wigglesworthia glossinidia (Yale colony)* | GCF_000247565.1 (26) | 134 | 349 | 824 | 608 |
| Phu-*Pediculus humanus corporis* holobiont | - | 575 | 2,329 | 3,760 | 2,132 |
| Phu-*Pediculus humanus* corporis USDA | GCA_000006295.1 (27) | 282 | 2,059 | 2,403 | 1,414 |
| Phu-*Candidatus* riesia pediculicola USDA | GCF_000093065.1 (27) | 105 | 287 | 786 | 494 |
| *Sitophilus oryzae* holobiont | - | 739 | 4,410 | 4,459 | 2,637 |
| *Sitophilus oryzae* Bouriz | GCF_002938485.1 (28) | 298 | 3,654 | 2,676 | 1,586 |
| *Candidatus* Sodalis pierantonius SOPE | GCF_000517405.1 (29) | 274 | 762 | 1,513 | 1,046 |

[a] http://www.whiteflygenomics.org/ftp/MEAM1/endosymbionts; [b] http://www.whiteflygenomics.org/ftp/MED/v1.0; [c] https://vectorbase.org/

**Table S2**. Organisms with GSMM available in the literature and used for our comparative analysis with ArtSymbioCyc reconstructions.

| **Organism** | **Short-name model (first author)** | **Reference** |
|---|---|---|
| *Acyrthosiphon pisum* (pea aphid) | Blow | (30) |
| *Buchnera aphidicola* of *A. pisum* | Blow | (30) |
| *Hamiltonella defensa* of *A. pisum* | Blow | (30) |
| *Bemisia tabaci* (whitefly) | Ankrah | (31) |
| *Portiera aleyrodidarum* of *B. tabaci* | Ankrah | (31) |
| *Hamiltonella defensa* of *A. pisum* | Ankrah | (31) |
| *Sodalis glossinidius* of *Glossina morsitans* | Belda | (32) |
| *Buchnera aphidicola* of *Cinara cedri* | Ponce-de-Leon | (33) |
| *Serratia symbiotica* of *Cinara cedri* | Ponce-de-Leon | (33) |
| *Drosophila melanogaster* (fruit fly) | Schönborn | (34) |
| *Drosophila melanogaster* (fruit fly) | Cesur | (35) |

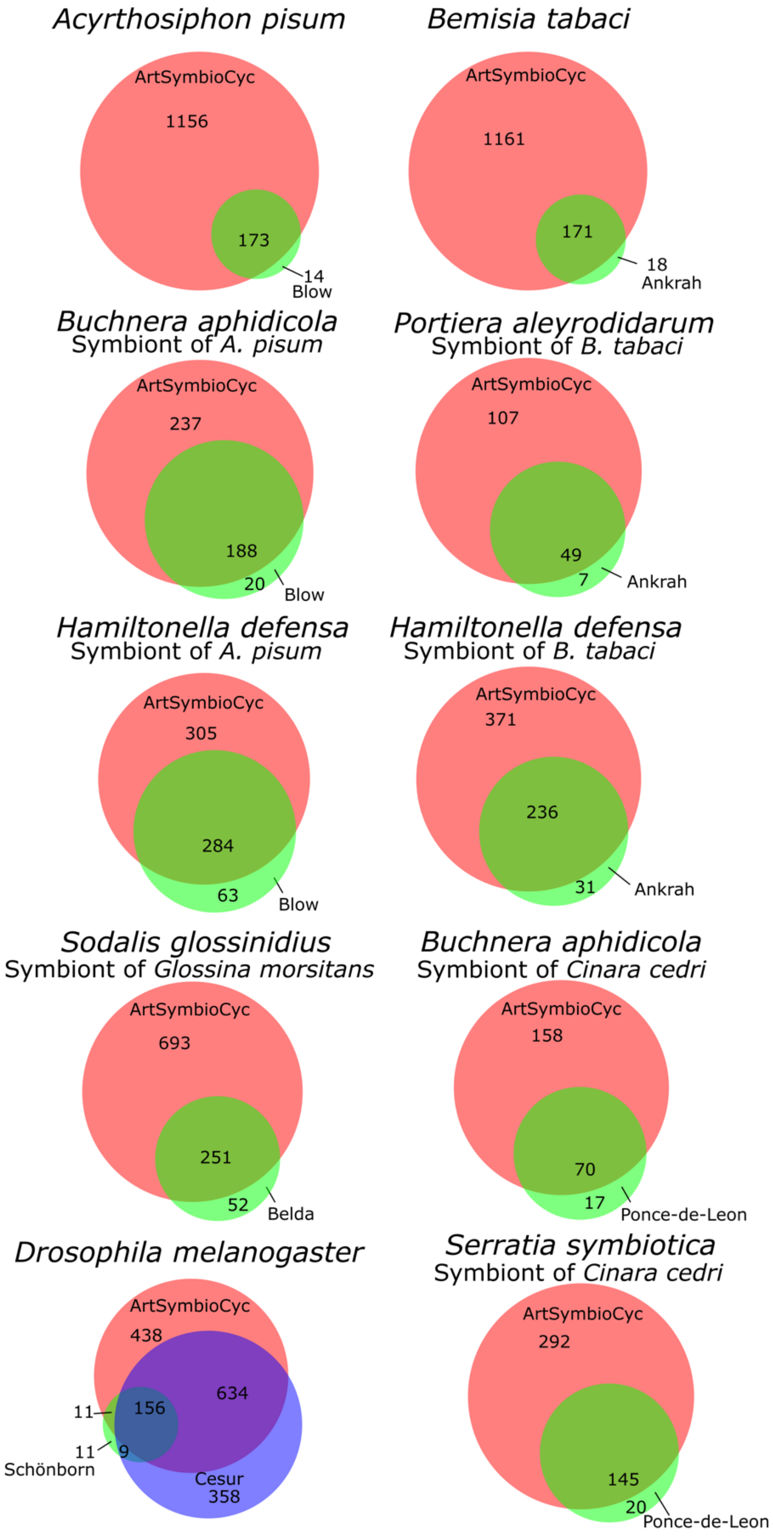


**Figure S1**. Venn diagrams for comparison of metabolic reconstructions of arthropods and their symbiotic bacteria. Red: metabolic reconstructions from the database ArtSymbioCyc, and in green or blue from GSMMs developed for FBA, with the first author of the associated article mentioned. See Table S2 for complete reference. EC numbers were used as a comparison metric, and the area is proportional to the number of EC numbers either specific or overlapping. Venn diagrams were made using the tool BioVenn https://www.biovenn.nl (36).

**Table S3**. Amino acids biosynthesis pathways comparison between the three aphid holobionts from the ArtSymbioCyc database collection: *Acyrthosiphon pisum* holobiont (Api-holobiont_bucap) with *Buchnera aphidicola* (Api-*B._aphidicola*_APS), *Cinara cedri* holobiont (Cce-holobiont) with *Buchnera aphidicola* (Cce-B_aphidicola_BCc) and *Serratia symbiotica* (Cce-S._symbiotica_Cc) and *Sipha maydis* holobiont (Sma-holobiont) with *Buchnera aphidicola* (Sma-B._aphidicola_Sm_Midelt) and *Serratia symbiotica* (Sma-S._symbiotica_Sm_Midelt). Green cells correspond to functional pathways automatically annotated and available to users from the interface. Red cells correspond to absences of automatically annotated pathways. Blue cells correspond to manually reconstructed pathways (the individual reactions are present but were not automatically gathered into pathways). Violet cells correspond to incomplete pathways manually tagged (pathways automatically annotated but with one or more lacking enzymes). The complete table can be directly visualized on the interface from this link.

| Amino acid | Pathways | *Api-B. aphidicola* APS | *Api-holobiont (bucap)* | *Cce-B. aphidicola* BCc | *Cce-S. symbiotica* Cc | *Cce-holobiont* | *Sma-B. aphidicola* Sm_Midelt | *Sma-S. symbiotica* Sm Midelt | *Sma-holobiont* |
|---|---|---|---|---|---|---|---|---|---|
| Gly | glycine biosynthesis I from serine | ✓ | ✓ | ✓ | ✓ | ✓ | ✓ | ✓ | ✓ |
| | glycine biosynthesis II (eukaryote glycine clivage complexe) | | ✓ | | | ✓ | | | ✓ |
| | glycine biosynthesis III from glyoxylate | | ✓ | | | ✓ | | | ✓ |
| | glycine biosynthesis IV from threonine | | ✓ | | | ✓ | | | ✓ |
| Ala | L-alanine biosynthesis II from pyruvate | | ✓ | | | ✓ | | | ✓ |
| | L-alanine biosynthesis III from Cysteine | ✓ | ✓ | ✓ | ✓ | ✓ | ✓ | ✓ | ✓ |
| Arg | L-arginine biosynthesis I (via L-ornithine) from glutamate | ✓ | n.f.[a] | from ornithine | | n.f. | from ornithine | | n.f. |
| Asn | L-asparagine biosynthesis I from aspartate | | ✓ | | | ✓ | | | ✓ |
| Asp | L-aspartate biosynthesis from oxaloacetate and glutamate | | ✓ | | | ✓ | | | ✓ |
| Cys | L-cysteine biosynthesis I from serine | ✓ | ✓ | | ✓ | ✓ | | | from HCYS[b] |
| Glu | L-glutamate biosynthesis I from glutamine | | ✓ | | | ✓ | | | ✓ |
| | L-glutamate biosynthesis II from 2-oxoglutarate | | ✓ | | | ✓ | | | ✓ |
| | L-glutamate biosynthesis IV from 2-oxoglutarate and gln | | ✓ | | | ✓ | | | ✓ |
| Gln | L-glutamine biosynthesis I | | ✓ | | | ✓ | | | ✓ |
| His | L-histidine biosynthesis from D-ribose-5-P | ✓ | ✓ | ✓ | | ✓ | ✓ | | ✓ |
| Ile | L-isoleucine biosynthesis I (from threonine) | | ✓ | | | ✓ | | | ✓ |
| Leu | L-leucine biosynthesis from 3-methyl-2-oxobutanoate | ✓ (incomplete)[c] | ✓ | ✓ (incomplete) | | ✓ | ✓ (incomplete) | | ✓ |
| Lys | L-lysine biosynthesis I from aspartate | ✓ | ✓ | ✓ | ✓ (incomplete) | ✓ | ✓ | ✓ (incomplete) | ✓ |
| Met | L-methionine biosynthesis from S-Methyl-L-Methionine | | ✓ | | | ✓ | | | ✓ |

| | | | | | | | | | |
|---|---|---|---|---|---|---|---|---|---|
| Phe | L-phenylalanine biosynthesis I from chorismate | √ (incomplete) | √ | √ (incomplete) | | √ | √ (incomplete) | | √ |
| | Chorismate biosynthesis from erythrose-4P | √ | √ | √ | | √ | √ | √ | √ |
| Pro | L-proline biosynthesis I (from L-glutamate) | | √ | | | √ | | | √ |
| Ser | L-serine biosynthesis I from 3P-D-glycerate | | √ | | | √ | | √ (incomplete) | √ |
| Thr | L-threonine biosynthesis from L-homoserine | √ | √ | √ | | √ | √ | | √ |
| | L-homoserine biosynthesis from aspartate | √ | √ | √ | | √ | √ | √ (incomplete) | √ |
| Trp | L-tryptophan biosynthesis from chorismate | √ | √ | | √ | √ | √ | | √ |
| | Chorismate biosynthesis from erythrose-4P | √ | √ | √ | | √ | √ | √ | √ |
| Tyr | L-tyrosine biosynthesis IV from phenylalanine | | √ | | | √ | | | √ |
| Val | L-valine biosynthesis from pyruvate | √ (incomplete) | √ | √ (incomplete) | | √ | √ (incomplete) | | √ |

[a] not found as an entire pathway per se but can be manually reconstructed (reactions are present); [b] homocysteine; [c] the pathway is incomplete (one or more enzymes are lacking).

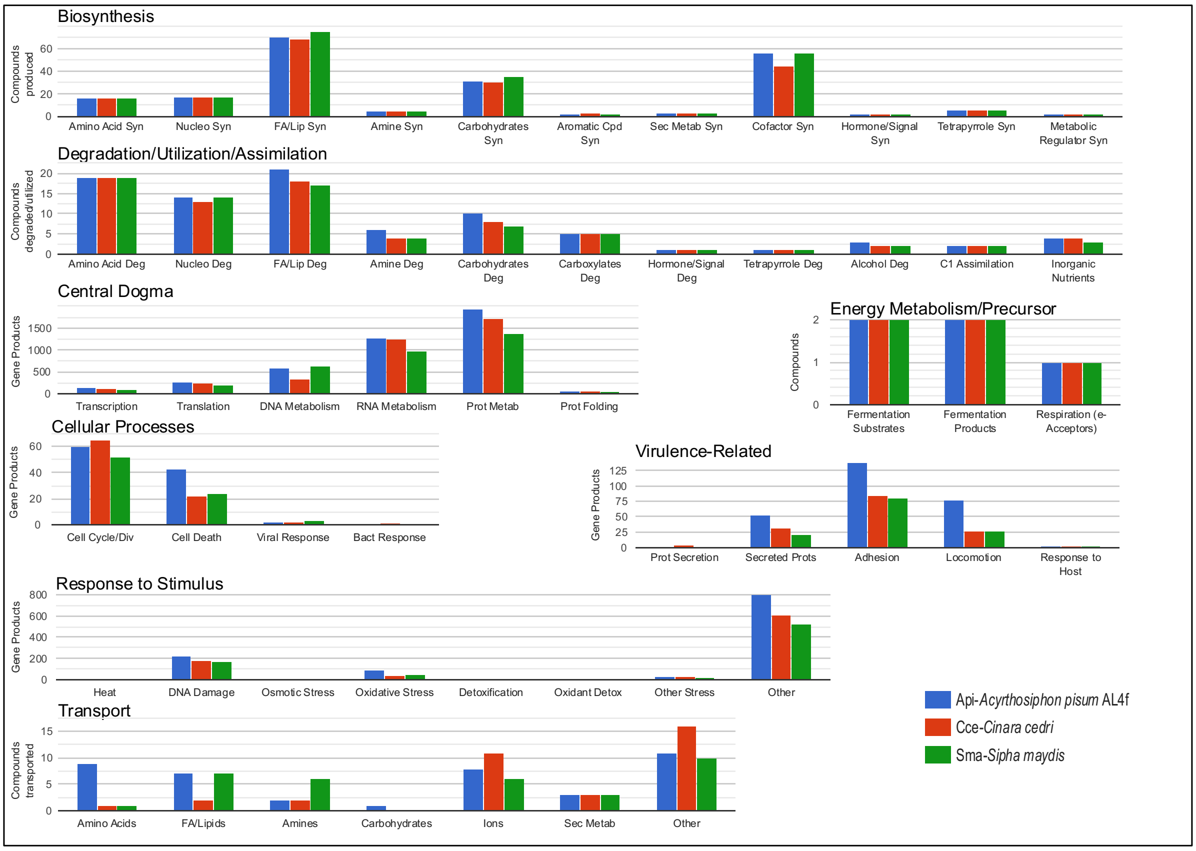


**Figure S2**. Central Metabolic pathway comparison between *A. pisum, C. cedri* and *S. maydis* using the Comparative Genome Dashboard (37). These comparisons can be visualized directly on the ArtSymbioCyc interface, and users can then click on each bar chart to see which pathways are specifically present or absent in the three organisms. As the plots are interactive, it is possible to access these pathways to determine the reactions and metabolites of which they are composed.
